# A Low-Resource Machine-Learning Framework for Cold-Stress Early Warning in Aquaculture Nursery Ponds Using Manual Temperature Readings

**DOI:** 10.64898/2026.08.01.742243

**Authors:** Imran Bin Younos, Nushrat Jahan

## Abstract

Cold stress is a recurring risk in tropical and subtropical aquaculture nursery ponds, yet warning tools remain limited where continuous automated sensors are impractical. This study developed a low-resource cold-stress early warning framework using four years (2022-2025) of 6-hourly manual air and pond-water temperature readings from a Nile tilapia (*Oreochromis niloticus*) nursery pond in Cumilla, Bangladesh. Models were fitted on 2022-2023, validated on 2024 for threshold selection, and tested on 2025 as an independent year. Cold stress (daily mean water temperature <20°C) occurred on 133 days; heat stress (>35°C) on only 4 days. Air-water coupling was strong overall (r = 0.976) but weakened in winter (r = 0.776) and further within the 18-22°C boundary zone where cold-stress classification is most sensitive. Solar radiation only marginally increased boundary-zone classification AUC from 0.782 to 0.789. In 6-hour regression, the same-hour-yesterday baseline (MAE = 1.117°C) nearly matched Extreme Gradient Boosting (XGBoost) with MAE of 1.116°C, cold-zone bias +0.36°C, and train-test gap 0.02°C; Random Forest (RF) and Long Short-Term Memory (LSTM) had MAEs of 1.195°C and 1.244°C, respectively. For cold-stress classification, Multiple Linear Regression (MLR) gave the highest F1 (0.755), while XGBoost provided the more protective operating point, detecting 95 of 108 cold-stress readings at 6-hour lead time (sensitivity = 0.880, F1 = 0.739). XGBoost warning skill extended to 12, 18, and 24-hour lead times, with F1 scores of 0.722, 0.646, and 0.704, respectively. The framework converts routine manual thermometer readings into short-lead cold-stress alerts for nursery management decisions. Multi-pond validation is needed before deployment.

## Introduction

Aquaculture is essential for supplying the world’s increasing demand for fish and other aquatic foods. Strengthening the sustainability and resilience of aquaculture systems is therefore a major priority of the Blue Transformation initiative led by the Food and Agriculture Organization (FAO, 2024). In Bangladesh, pond aquaculture is a cornerstone of fish production, and Nile tilapia (*Oreochromis niloticus*) has become one of the most widely cultured species because of its rapid growth, high productivity, broad adaptability, and compatibility with diverse culture systems (Rahman et al., 2021). Despite its economic importance, pond aquaculture in Bangladesh remains exposed to weather variability, and climate-service studies have identified temperature and rainfall variability as operational risks requiring species-specific advisories (Hossain et al., 2021; Montes et al., 2022).

Temperature is one of the most influential environmental factors governing fish physiology, metabolism, growth, and reproduction in nursery and hatchery systems, particularly in regions where seasonal water temperature fluctuations are pronounced (Azaza et al., 2008; Leonard and Skov, 2022; Nivelle et al., 2019; Nobrega et al., 2020). Among warm-water aquaculture species, Nile tilapia is especially relevant because of its economic importance and well-documented thermal sensitivity. Although tilapia exhibit a relatively broad thermal tolerance, their growth performance and reproductive success are highly dependent on maintaining suitable temperature conditions. Thermal requirements vary among developmental stages, reflecting differences in temporal and spatial distribution patterns during early life-history development (Abd El-Hack et al., 2022). In Nile tilapia, fingerlings perform best within 25 to 30°C, while juveniles require slightly warmer conditions of 27 to 32°C, with a lower lethal threshold near 11 to 12°C and an upper lethal limit approaching 42°C (Leonard and Skov, 2022; Nivelle et al., 2019). At the upper thermal extreme, temperatures exceeding 35–37°C elicit heat-shock protein responses, suppress growth-related gene expression, and cause histopathological changes to gill lamellae in juvenile and fingerling tilapia (Chakrabarty et al., 2025; Islam et al., 2022). For nursery and hatchery operations, however, the relevant management concern extends well below the point of lethal exposure. Sublethal cold conditions suppress feeding, growth, movement, and recovery in fry and fingerlings (Nobrega et al., 2020). Since optimal growth and feed conversion in these early life stages requires temperatures between 26 and 30°C (Azaza et al., 2008; Nobrega et al., 2020), both cold and heat stress represent operationally relevant hazards.

Temperature monitoring in low-resource hatchery settings is often manual. Smart aquaculture studies increasingly use IoT networks, cloud dashboards, and continuous sensors (Zhao et al., 2021), and fishpond cold-snap systems combine weather models with pond sensors (Li et al., 2024). These systems are valuable but do not match all farm contexts. A practical Bangladesh nursery warning tool should work from thermometer readings that farmers can obtain without continuous automated sensor hardware. This constraint is particularly relevant for smallholder operations where capital investment in sensor infrastructure remains limited (Hossain et al., 2021).

Air temperature is an attractive predictor because it is cheap to observe and physically connected to surface-water heat exchange. In hydrology, nonlinear air-water relationships and thermal hysteresis have been studied extensively (Caissie, 2006; Mohseni et al., 1998). In aquaculture, machine-learning models have been applied to predict water-quality variables from manually collected observations (Zambrano et al., 2021). Siddique et al. (2024) modelled water temperature in a tilapia broodfish pond in Mymensingh using ARIMAX and identified air temperature and solar intensity as important exogenous factors. Resnick et al. (2024) used an energy-balance model to simulate pond water temperature at daily and seasonal timescales in southwest and northeast Bangladesh using publicly available weather data. Together these studies show that air temperature tracks pond water temperature well in aquaculture ponds. What matters for a warning system is where that tracking weakens, and the sharpest case is near the cold-stress threshold, where a small prediction error flips the classification. The present study advances this literature by using six hourly observations from a nursery pond, separating regression accuracy from warning performance, reporting precision, recall, and false alarms for cold-stress decisions, and testing the final warning framework on an independent year.

This study uses a four-year empirical pond record from a Nile tilapia nursery to move from temperature prediction to risk decision. A first step is to determine which thermal hazard actually dominated this pond: cold stress or heat stress. The analysis then asks how reliably air temperature tracks pond water temperature across seasons, and where that coupling loosens in the 18–22°C range where classification errors matter most. Solar radiation is tested as a possible supplementary predictor in that near-threshold zone. The modelling component then compares Multiple Linear Regression (MLR), Random Forest (RF), Extreme Gradient Boosting (XGBoost), and Long Short-Term Memory (LSTM) within a 6-hour-ahead early-warning framework, evaluated on whether warnings would have been timely and accurate in the independent 2025 year. The operating constraint is the absence of continuous automated sensing. The proposed solution is a threshold-based warning built from routine manual readings, evaluated through an independent-year test of warning skill against false-alarm burden.

We tested four hypotheses: (1) that cold stress dominates heat stress as the primary thermal hazard and varies across years; (2) that solar radiation is associated with cold-stress status within the boundary zone; (3) that nonlinear or sequence-aware models reduce cold-zone prediction bias relative to MLR; and (4) that a 6-hour thermometer-based warning framework provides meaningful operational warning in an independent test year. These hypotheses directly address the lack of year-scale thermal hazard comparisons, the uncertainty about solar radiation’s practical contribution, the unclear advantage of nonlinear models for cold-zone prediction, and the absence of tested low-cost, practical warning systems for small-scale aquaculture facilities where IoT infrastructure is unavailable.

## Methods

### Study site and measurements

The study was conducted in a Nile tilapia (*Oreochromis niloticus*) nursery pond in Cumilla, Bangladesh. The rectangular pond measured 40 m × 62.5 m, with a surface area of 2,500 m^2^ (0.25 ha). The operational water depth ranged from 1.52 to 1.68 m, giving an approximate water volume of 4,000 m^3^. Air and pond-water temperature were recorded at 6-h intervals (06:00, 12:00, 18:00, 00:00) from 1 January 2022 to 31 December 2025. Air temperature was measured under shaded conditions at the pond bank. Water temperature was measured at a fixed near-surface location (15-20 cm depth), consistent with standard pond monitoring protocols (Boyd and Tucker, 1998). All field records were obtained using a calibrated glass thermometer with a graduation resolution of 0.5°C, which represents the measurement uncertainty for the recorded dataset.

### Data preparation

The curated dataset comprised 1,461 daily records (5,844 total observations) with four air-temperature and four water-temperature readings per day. Before analysis the record was screened for transcription slips and physically implausible spikes; a small number of obvious single-point recording errors were corrected against the adjacent readings on the same day, and no other values were changed. Missing temperature records (under 1% of total observations) were then filled by linear interpolation prior to model training. Daily mean temperature (Tmean) and diurnal temperature range (DTR) were computed as:

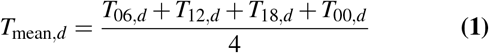

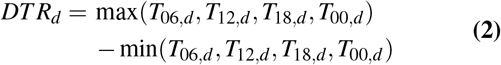

### Stress definitions and daily analyses

Seasons followed the Bangladesh Meteorological Department convention: winter (December-February), pre-monsoon (March-May), monsoon (June-September), and post-monsoon (October-November). Daily cold stress was defined as mean pond-water temperature <20°C, the threshold below which feeding and growth are substantially reduced in juvenile Nile tilapia (Charo-Karisa et al., 2005). Daily heat stress was defined as mean pond-water temperature >35°C. Days with mean water temperature <18°C and days with minimum water temperature <14°C were recorded as severity indicators. Consecutive cold-stress days were grouped as events.

Air-water thermal coupling was assessed with Pearson correlation and ordinary least squares (OLS) regression between daily mean air and water temperature for (i) the full record, (ii) by season, and (iii) within the 18-22°C air-temperature boundary zone, where cold-stress classification is most sensitive to prediction error. The 18–22°C subset was used as a decision-relevant descriptive window, not as a physical break-point.

Daily solar radiation and rainfall data were obtained from the NASA Prediction of Worldwide Energy Resources (POWER) Release 9 (Stackhouse et al., 2021) via the NASA POWER API (https://power.larc.nasa.gov/) for the grid cell covering the study site in Cumilla, Bangladesh (meteorological resolution 0.5°× 0.625°, radiation resolution 1° × 1°). The variables retrieved were all-sky surface shortwave downward irradiance (ALLSKY_SFC_SW_DWN, converted to MJ m^−2^ d^−1^) and bias-corrected precipitation (PRECTOTCORR, mm d^−1^). These data were treated as supporting regional estimates, not pond-level measurements. A full-range OLS model tested the association of solar radiation and rainfall with daily mean water temperature after accounting for air temperature. A boundary-zone analysis then compared solar radiation on cold-stress versus non-cold-stress days using the Mann-Whitney U test (Mann and Whitney, 1947), Cohen’s d effect size (Cohen, 1988), and logistic regression area under the receiver operating characteristic curve (AUC).

### Six-hour forecasting dataset

The four daily observations were arranged as a continuous 6-h sequence (06:00, 12:00, 18:00, 00:00). Each row represented a decision time i, with the target being the pond-water temperature at the next observation (6-h horizon):

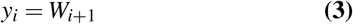

where W denotes pond-water temperature. The predictor vector comprised autoregressive air- and water-temperature lags, calendar indicators, and derived thermal features:

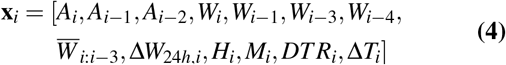

where A is air temperature,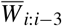 is the rolling mean of the four most recent water readings, Δ*W*_24*h,i*_ = *W*_*i*_ − *W*_*i* − 4_ is the 24-h water-temperature change, H is decision hour, M is month, DTR is diurnal temperature range, and ΔT is the current air-water temperature difference. The term *W*_*i* − 3_ represents the same-target-hour water temperature from the previous day. Future water temperature and NASA POWER variables were not used as inputs. Two naive baselines were evaluated alongside trained models: (1) persistence (*ŷ*_*i*_ = *W*_*i*_) and (2) same-hour-yesterday. (*ŷ*_*i*_ = *W*_*i*−3_) Rows lacking sufficient lag history or a next-step target were excluded.

### Model development

Training, validation, and testing followed a chronological (walk-forward) design to preserve temporal order and prevent data leakage (Bergmeir and Benítez, 2012; Tashman, 2000). Target readings from 2022-2023 were used for model fitting; readings from 2024 were held out for validation, early stopping, and alert-threshold selection; all readings from 2025 were reserved as the independent test year. Data were not shuffled. Four regression models were compared. Multiple Linear Regression (MLR) served as the linear baseline:

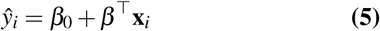

Random Forest (RF) was used as a bagged ensemble of regression trees (Breiman, 2001). Extreme Gradient Boosting (XGBoost) was used as a regularized gradient-boosted tree model (Chen and Guestrin, 2016). Long Short-Term Memory (LSTM) was used as a recurrent neural network for ordered 6-h thermal sequences (Hochreiter and Schmidhuber, 1997). RF hyperparameters were tuned via five-fold time-series cross-validation within the fitting period (Bergmeir et al., 2018), searching: *n*_estimators_ ∈ {100, 200, 500}, max depth ∈ {unrestricted, 10, 20}, and min samples leaf ∈ {1, 2, 4}. The selected configuration used 100 trees, maxdepth =20, and minsample_leaf =4. XGBoost was tuned with the same validation strategy, searching: *n*_estimators_ ∈ {100, 200, 300}, learning rate ∈ {0.01, 0.05, 0.10}, max depth ∈ {3, 5, 6,} and subsample ∈ {0.8, 1.0}. The selected configuration used 200 trees, learning_rate = 0.05, maxdepth = 3, and subsample = 0.8. MLR inputs were z-score standardized using fitting-period statistics.

The LSTM architecture comprised two recurrent layers (64 and 32 units), each followed by dropout (rate = 0.2), and a single-neuron dense output layer. It was trained with the Adam optimizer (Kingma and Ba, 2015) using learning_rate = 0.001, batch_size = 64, and MSE loss for up to 100 epochs, with early stopping after 10 epochs without validation-loss improvement. LSTM inputs were standardized using training-set parameters only.

### Evaluation and early-warning assessment

Regression performance was evaluated on the 2025 test year using mean absolute error (MAE), root mean squared error (RMSE), coefficient of determination (R^2^), and bias (Willmott, 1981):

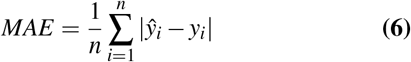

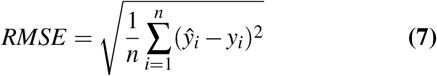

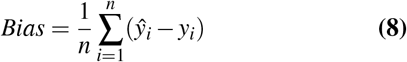

Metrics were computed for the full test year and for the cold subset (observed water temperature <20°C). Train-test MAE gap was calculated as test MAE minus train MAE. For early-warning conversion, the continuous 6-h prediction was thresholded into a binary cold-stress alert. For model m, an alert was issued when 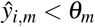, where threshold *θ*_*m*_ was selected on the 2024 validation set. Candidate thresholds from 16.0 to 24.0°C were evaluated at 0.1°C increments; the threshold maximizing validation F1 score was selected, with sensitivity, specificity, and proximity to the 20°C biological threshold used as tie-breakers. Because cold-stress readings were the minority class, precision and F1 were interpreted alongside ROC AUC (Saito and Rehmsmeier, 2015). Test-year warning performance was reported as sensitivity, specificity, precision, F1 score, ROC AUC, and false alarms per true warning.

Uncertainty in warning metrics was quantified using 1,000 moving-block bootstrap resamples (Künsch, 1989; Lahiri, 2003) of the 2025 chronological test sequence. Each resample comprised contiguous blocks of 28 readings (one week of 6-h observations), resampled with replacement to the original test-set length. Ninety-five percent confidence intervals were derived from the 2.5th and 97.5th percentiles of the bootstrap distribution.

Lead-time sensitivity was assessed by reconstructing the prediction target at 12, 18, and 24 h (two, three, and four 6-h intervals ahead). For each horizon, predictor variables were restricted to measurements available at or before the decision timestamp. Separate XGBoost models were fitted for each horizon using the same chronological training procedure, hyperparameter grid, and threshold-selection method.

### Software

Analyses were performed in Python 3.11 using pandas 2.1, NumPy 1.26, SciPy 1.11, statsmodels 0.14, scikit-learn 1.3, XGBoost 2.0, and TensorFlow/Keras 2.15. Random seeds were fixed at 42 for Python, NumPy, and TensorFlow to ensure reproducibility.

## Results

### Thermal conditions and seasonal patterns

Daily mean pond-water temperature was 27.61 ± 4.68°C (range: 15.75-35.50°C; Table 1). Daily mean air temperature was 27.45 ± 4.95°C (range: 15.50-37.25°C). Mean DTR was lower for water (5.96°C) than for air (8.66°C), indicating thermal damping by the pond. The four-year time series is shown in Figure 1. Winter was the coolest season in every year. Mean winter water temperature was 20.80, 20.49, 20.06, and 21.43°C in 2022, 2023, 2024, and 2025, respectively. Monsoon means ranged from 29.82 to 31.85°C. Monthly distributions are shown in Figure S1, and seasonal diurnal profiles are shown in Figure 2.

**Table 1.** Descriptive statistics of daily air and pond water temperature and thermal stress counts by year and season (2022–2025).

| Year | Season | n | Water mean (°C) | Water SD (°C) | Water min (°C) | Water max (°C) | Water DTR (°C) | Air mean (°C) | Air DTR (°C) | Water:air DTR ratio | Cold-stress days | Heat-stress days |
| --- | --- | --- | --- | --- | --- | --- | --- | --- | --- | --- | --- | --- |
| 2022 | Winter | 90 | 20.80 | 1.508 | 16.25 | 25.88 | 10.22 | 20.38 | 12.72 | 0.803 | 23 | 0 |
|  | Pre-monsoon | 92 | 29.48 | 2.466 | 21.50 | 32.00 | 3.076 | 28.58 | 8.125 | 0.379 | 0 | 0 |
|  | Monsoon | 122 | 30.92 | 0.957 | 28.25 | 33.00 | 2.705 | 30.11 | 6.668 | 0.406 | 0 | 0 |
|  | Post-monsoon | 61 | 27.55 | 1.747 | 24.50 | 30.75 | 4.631 | 26.93 | 10.21 | 0.454 | 0 | 0 |
| 2023 | Winter | 90 | 20.49 | 1.786 | 16.38 | 24.00 | 9.367 | 20.16 | 11.72 | 0.799 | 35 | 0 |
|  | Pre-monsoon | 92 | 30.78 | 3.179 | 25.50 | 35.00 | 5.141 | 31.56 | 7.457 | 0.690 | 0 | 0 |
|  | Monsoon | 122 | 31.85 | 1.923 | 27.50 | 35.50 | 3.877 | 31.92 | 4.713 | 0.823 | 0 | 3 |
|  | Post-monsoon | 61 | 27.40 | 3.039 | 19.00 | 31.75 | 4.820 | 27.36 | 8.164 | 0.590 | 2 | 0 |
| 2024 | Winter | 91 | 20.06 | 2.225 | 15.75 | 24.75 | 8.824 | 19.50 | 11.70 | 0.754 | 49 | 0 |
|  | Pre-monsoon | 92 | 30.00 | 2.457 | 22.00 | 35.25 | 6.217 | 30.04 | 9.554 | 0.651 | 0 | 1 |
|  | Monsoon | 122 | 29.82 | 1.308 | 27.00 | 33.50 | 3.836 | 30.13 | 5.246 | 0.731 | 0 | 0 |
|  | Post-monsoon | 61 | 25.82 | 3.283 | 19.00 | 30.25 | 4.984 | 26.11 | 7.049 | 0.707 | 6 | 0 |
| 2025 | Winter | 90 | 21.43 | 1.661 | 17.75 | 24.75 | 8.067 | 21.19 | 10.34 | 0.780 | 18 | 0 |
|  | Pre-monsoon | 92 | 29.23 | 1.464 | 26.00 | 31.75 | 10.11 | 29.41 | 12.40 | 0.815 | 0 | 0 |
|  | Monsoon | 122 | 31.64 | 1.247 | 28.75 | 34.00 | 4.779 | 31.47 | 7.811 | 0.612 | 0 | 0 |
|  | Post-monsoon | 61 | 29.42 | 2.374 | 23.75 | 33.00 | 6.869 | 29.18 | 7.607 | 0.903 | 0 | 0 |

DTR = diurnal temperature range. Cold stress: daily mean water temperature <20°C; heat stress: >35°C. Seasons follow Bangladesh Meteorological Department convention: winter (Dec–Feb), pre-monsoon (Mar–May), monsoon (Jun–Sep), post-monsoon (Oct–Nov). Year shown for first season of each year only.

**Figure 1.**
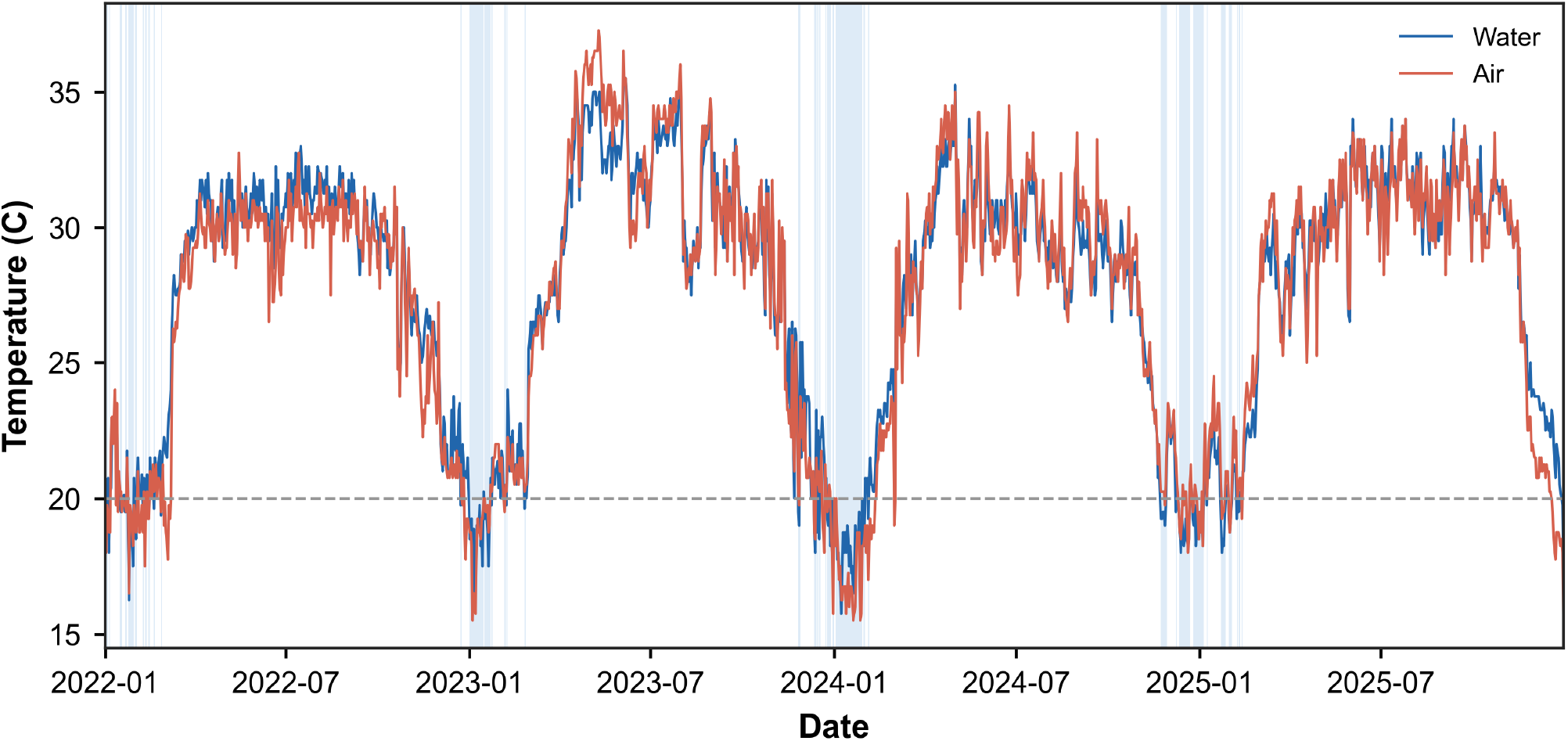
Daily mean air and pond water temperature at the Nile tilapia (*Oreochromis niloticus*) nursery pond in Bangladesh (January 2022 to December 2025). Shaded intervals indicate periods of cold stress (daily mean water temperature below 20°C).

**Figure 2.**
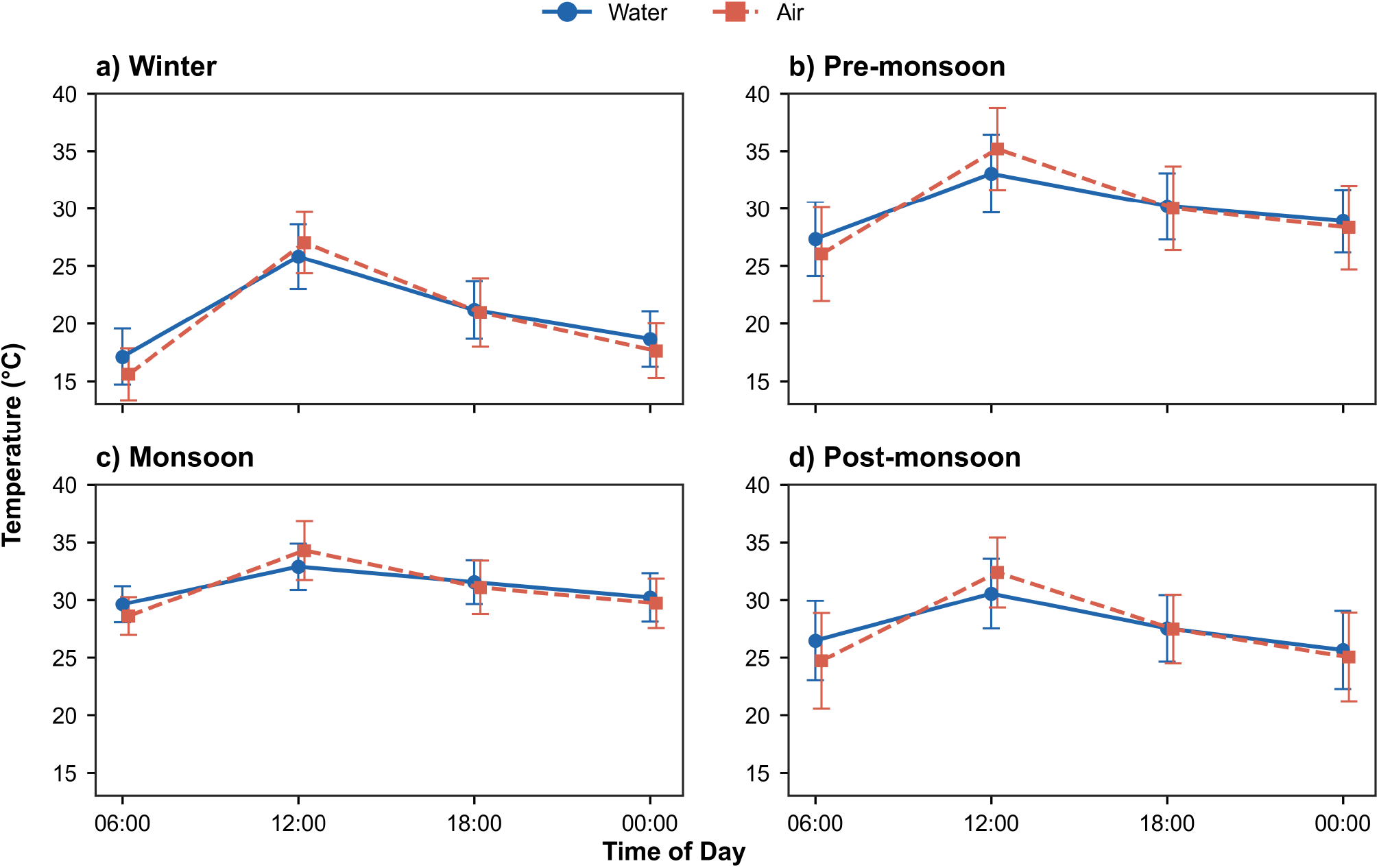
Mean diurnal profiles of air and pond water temperature at the four daily observation times (06:00, 12:00, 18:00, 00:00) for each season: (a) winter, (b) pre-monsoon, (c) monsoon, (d) post-monsoon. Season boundaries follow the Bangladesh Meteorological Department classification.

### Cold-stress burden and interannual variation

Cold stress occurred on 133 days, compared with 4 heat-stress days. Annual cold-stress counts were 23, 37, 55, and 18 days in 2022-2025, respectively (Table 2). The highest annual burden occurred in 2024, with 55 cold-stress days and a winter air mean of 19.5°C. In 2025, cold stress declined to 18 days while winter air temperature averaged 21.2°C. Cold-stress events were longest in 2024. That year had 7 events (mean duration 7.86 d; maximum 27 d), compared with 8 events in 2025 (mean duration 2.25 d; maximum 5 d). Days with mean water temperature below 18°C numbered 2, 6, 12, and 1 in 2022-2025, respectively. Cold-stress days were concentrated in January and December (Figure 3). January accounted for 14, 21, 29, and 12 cold-stress days in 2022-2025. December accounted for 11 days in 2023 and 19 days in 2024.

**Table 2.** Annual cold-stress event characteristics and thermal severity indicators for the nursery pond (2022–2025).

| Year | Cold-stress events | Total cold-stress days | Single-day events | Events >7 d | Mean duration (d) | Max duration (d) | Days mean <18°C | Days min <14°C | Heat-stress days | Winter air mean (°C) |
| --- | --- | --- | --- | --- | --- | --- | --- | --- | --- | --- |
| 2022 | 12 | 23 | 5 | 0 | 1.917 | 6 | 2 | 13 | 0 | 20.38 |
| 2023 | 13 | 37a | 7 | 1 | 2.846 | 14 | 6 | 5 | 3 | 20.16 |
| 2024 | 7 | 55a | 2 | 2 | 7.857 | 27 | 12 | 7 | 1 | 19.50 |
| 2025 | 8 | 18 | 4 | 0 | 2.250 | 5 | 1 | 0 | 0 | 21.19 |

<sup>a</sup> Totals include cold-stress days outside the winter season: 2 post-monsoon days in 2023 and 6 post-monsoon days in 2024. Cold stress: daily mean water temperature <20°C; heat stress: >35°C. Winter air mean computed over December to February.

**Figure 3.**
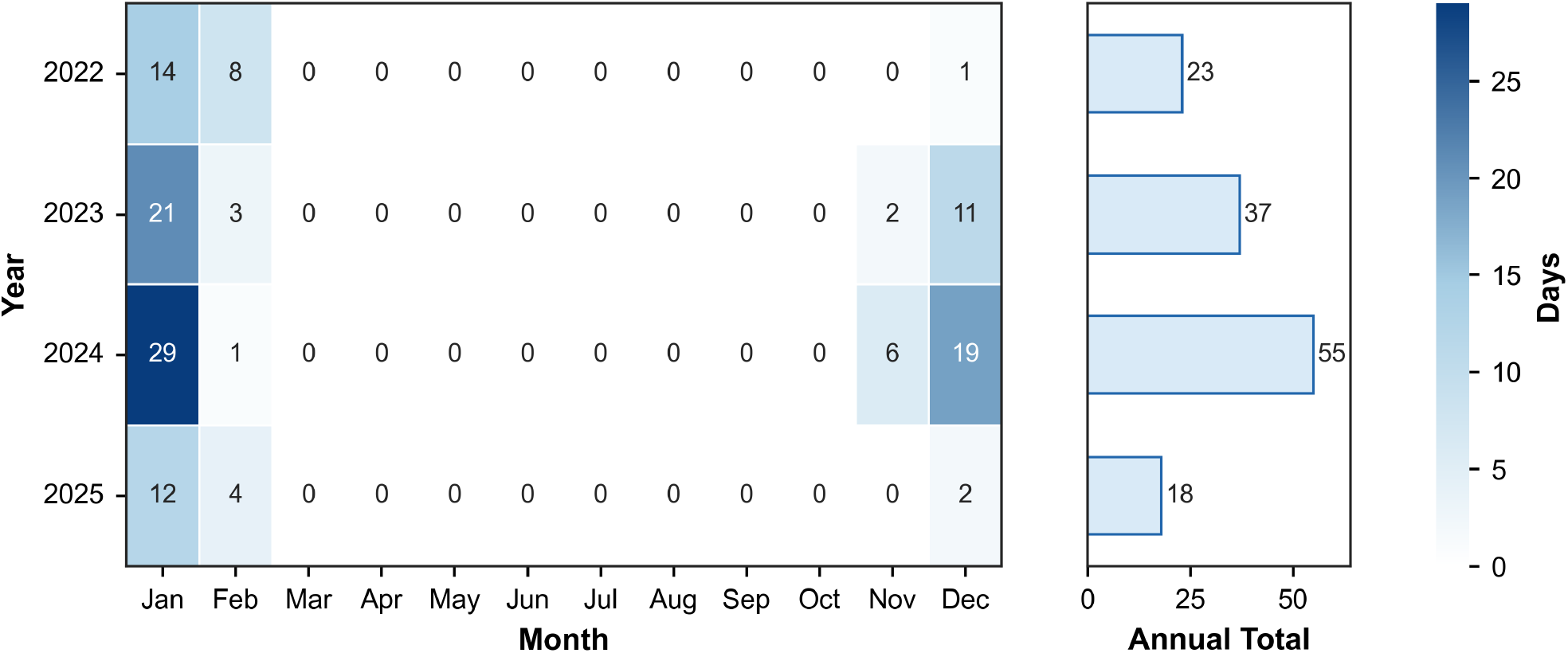
Monthly cold-stress day counts by year (2022–2025) with annual totals. Cold stress is defined as daily mean pond water temperature below 20°C. Cell values in the heat map show the number of cold-stress days per month-year combination.

### Air-water coupling and near-threshold variability

Daily mean air and water temperature were strongly correlated over the full record (r = 0.976). The full-record OLS slope was 0.922, with an intercept of 2.32°C and R^2^ = 0.952. Winter had the weakest seasonal coupling (r = 0.776, R^2^ = 0.603). Coupling was also weaker within the 18-22°C air-temperature boundary zone (n = 295; r = 0.541; R^2^ = 0.292), where cold-stress classification is most sensitive to small water-temperature errors. The boundary-zone slope (0.775) remained close to the full-record slope (0.922). Figure 4 shows the season-coloured air-water relationship and full-record linear fit, with 20°C retained only as a cold-stress decision reference.

**Figure 4.**
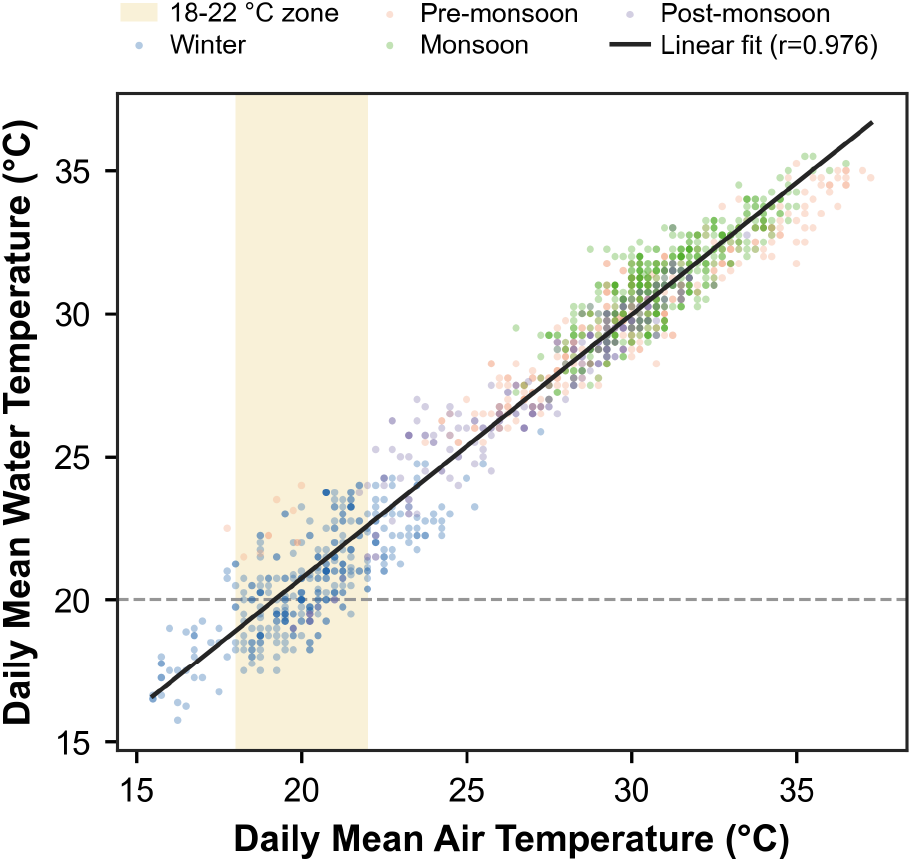
Daily mean air versus pond water temperature across the full four-year record (n = 1,461 days), colour-coded by season. The solid line is the full-record ordinary least squares fit (r = 0.976, R^2^ = 0.952). The shaded vertical band marks the 18–22°C air-temperature boundary zone; the dashed horizontal line indicates the 20°C cold-stress threshold.

### Solar radiation in the boundary zone

In the full annual OLS model, solar radiation was not significant after accounting for daily mean air temperature and rainfall (coefficient = 0.00521, p = 0.503; model R^2^ = 0.952). Rainfall had a small positive coefficient (0.01061, p = 0.00001).

Within the 18-22°C air-temperature boundary zone (n = 295; 103 cold-stress days and 192 non-cold days), mean solar radiation was 14.18 MJ m^−2^ d^−1^ on cold-stress days and 15.54 MJ m^−2^ d^−1^ on non-cold days (Figure 5). The difference was significant by Mann-Whitney U test (U = 7405.5, p < 0.001; Cohen’s d = -0.39). AUC changed from 0.782 with air temperature alone to 0.789 with air temperature plus solar radiation.

**Figure 5.**
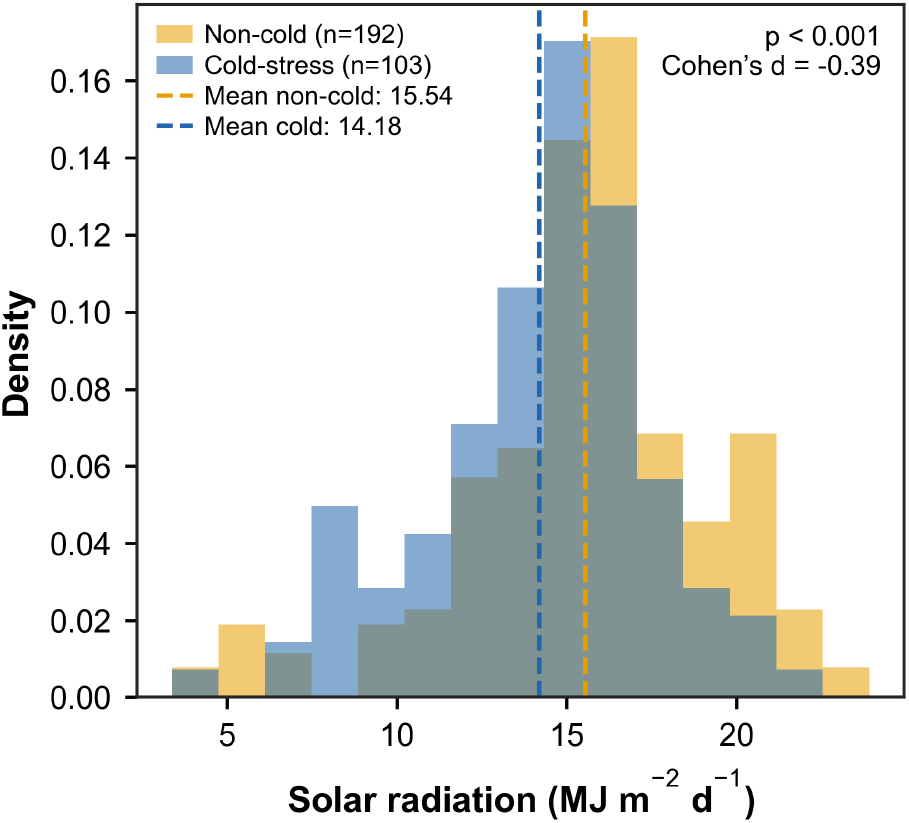
Distributions of daily solar radiation (MJ m^−^2 d^−^1 ) on cold-stress (n = 103) and non-cold (n = 192) days within the 18–22°C air-temperature boundary zone. Vertical dashed lines indicate group means. The Mann-Whitney U test result and Cohen’s d effect size are shown within the panel.

### Model performance and cold-zone residuals

On the independent 2025 sub-daily test set, the same-hour-yesterday baseline was strong, with MAE = 1.117°C, RMSE = 1.448°C and R^2^ = 0.924 (Table 3). XGBoost was the best trained regression model by a very small margin, with MAE = 1.116°C, RMSE = 1.427°C and R^2^ = 0.926. MLR had MAE = 1.135°C, RF had MAE = 1.195°C and LSTM had MAE = 1.244°C. True persistence from the decision-time water reading was much weaker (MAE = 3.658°C). The useful naive signal therefore came from same-target-hour thermal memory, not from one-step persistence.

**Table 3.** Six-hour-ahead regression performance of trained models and naive baselines on the independent 2025 test year, reported for the full test set and the cold subset.

| Model | Train MAE (°C) | Test MAE (°C) | Train RMSE (°C) | Test RMSE (°C) | Train R<sup>2</sup> | Test R<sup>2</sup> | MAE gap (°C) | Cold subset (n) | Cold MAE (°C) | Cold RMSE (°C) | Cold R<sup>2</sup> | Cold bias (°C) |
| --- | --- | --- | --- | --- | --- | --- | --- | --- | --- | --- | --- | --- |
| Persistence | — | 3.658 | — | 4.738 | — | 0.183 | — | 108 | 2.694 | 3.248 | -5.387 | 2.620 |
| SHY<sup>a</sup> | — | 1.117 | — | 1.448 | — | 0.924 | — | 108 | 1.037 | 1.232 | 0.080 | 0.370 |
| MLR | 1.316 | 1.135 | 1.809 | 1.453 | 0.887 | 0.923 | -0.182 | 108 | 1.312 | 1.601 | -0.553 | 1.163 |
| RF | 0.774 | 1.195 | 1.106 | 1.536 | 0.958 | 0.914 | 0.421 | 108 | 1.044 | 1.324 | -0.062 | 0.297 |
| XGBoost | 1.097 | 1.116 | 1.475 | 1.427 | 0.925 | 0.926 | 0.019 | 108 | 1.004 | 1.312 | -0.042 | 0.360 |
| LSTM | 1.281 | 1.244 | 1.701 | 1.632 | 0.900 | 0.903 | -0.037 | 108 | 1.265 | 1.776 | -0.910 | 1.066 |

MAE = mean absolute error; RMSE = root mean square error. MAE gap = test MAE - train MAE. <sup>a</sup> SHY = same-hour-yesterday naive baseline.

The train-test MAE gap was 0.02°C for XGBoost, 0.42°C for RF, -0.04°C for LSTM and -0.18°C for MLR. The RF observed-predicted six-hour time series is provided as Figure S2, and observed-predicted scatter diagnostics for all trained models are provided in Figure S3.

The 2025 cold subset contained 108 target readings below 20°C. Mean cold-zone bias was +1.16°C for MLR, +0.30°C for RF, +0.36°C for XGBoost and +1.07°C for LSTM (Figure 6). The same-hour-yesterday baseline had cold-zone bias of +0.37°C. Cold-subset R^2^ values were close to zero or negative for the trained models. The meaningful cold-range diagnostics are absolute error and bias, which stayed low for the tree models (XGBoost cold MAE = 1.004°C, cold bias = +0.36°C; Table 3).

**Figure 6.**
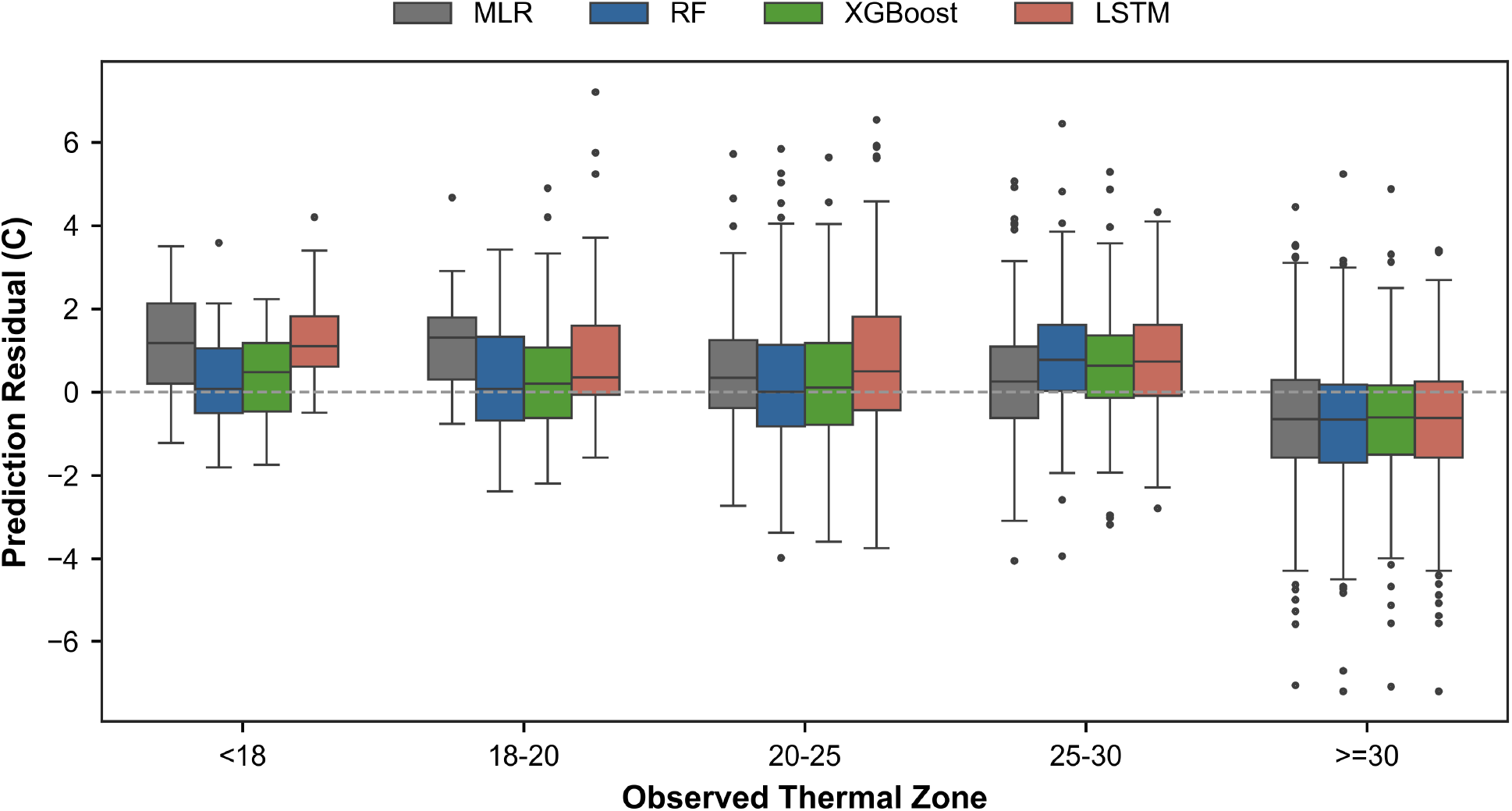
Prediction residuals (predicted minus observed pond water temperature, °C) across five observed temperature zones for each model on the 2025 test set. The dashed line at zero indicates unbiased prediction; positive values reflect systematic overestimation of water temperature.

Seasonal test errors are reported in Table S1. XGBoost MAE was 1.02°C in winter, 1.29°C in pre-monsoon, 1.09°C in monsoon and 1.03°C in post-monsoon. MLR MAE was 1.05, 1.27, 1.17 and 0.99°C for the same seasons. Time-of-day EWS results are reported in Table S2, and full coupling statistics are reported in Table S3.

### Six-hour early-warning performance

The 2025 EWS test set contained 1,460 target readings, including 108 cold-stress readings. Validation-selected thresholds clustered near 20°C for most models, ranging from 19.6°C for RF to 21.1°C for persistence (Table 4). ROC AUC was high across all trained models (Figure 7). The same-hour-yesterday baseline detected 103 of 108 cold-stress readings but produced 74 false alarms, giving sensitivity = 95.4%, precision = 58.2% and F1 = 72.3%. Among trained models, MLR produced the highest balanced warning score, with F1 = 75.5%, precision = 76.9% and 24 false alarms, but sensitivity was lower at 74.1%. XG-Boost gave the more protective operating point: 95 of 108 cold-stress readings were detected, with 13 missed events and 54 false alarms. Its sensitivity, specificity, precision and F1 were 88.0%, 96.0%, 63.8% and 73.9%, respectively. RF and LSTM gave F1 scores of 73.5% and 72.0%. Across the full independent 2025 test year, XGBoost alerts were concentrated during the cool-season periods, while most warm-season observations were correctly classified as non-cold (Figure 8).

**Table 4.** Six-hour-ahead cold-stress warning performance on 2025 sub-daily target readings. Confidence intervals are block-bootstrap 95% intervals from 1000 chronological resamples.

| Model | Trigger (°C) | TP | FN | FP | TN | Sensitivity | Sensitivity 95% CI | Specificity | Specificity 95% CI | Precision | Precision 95% CI | F1 | F1 95% CI | False alarms per true warning | ROC AUC |
| --- | --- | --- | --- | --- | --- | --- | --- | --- | --- | --- | --- | --- | --- | --- | --- |
| Persistence | 21.10 | 87 | 21 | 151 | 1201 | 0.806 | 0.691–0.867 | 0.888 | 0.832–0.934 | 0.366 | 0.241–0.466 | 0.503 | 0.356–0.603 | 1.736 | 0.914 |
| SHY<sup>a</sup> | 20.10 | 103 | 5 | 74 | 1278 | 0.954 | 0.902–0.988 | 0.945 | 0.916–0.968 | 0.582 | 0.456–0.678 | 0.723 | 0.609–0.802 | 0.718 | 0.986 |
| MLR | 20.10 | 80 | 28 | 24 | 1328 | 0.741 | 0.576–0.844 | 0.982 | 0.971–0.992 | 0.769 | 0.627–0.851 | 0.755 | 0.610–0.838 | 0.300 | 0.986 |
| RF | 19.60 | 86 | 22 | 40 | 1312 | 0.796 | 0.652–0.876 | 0.970 | 0.951–0.985 | 0.683 | 0.546–0.784 | 0.735 | 0.603–0.812 | 0.465 | 0.983 |
| XGBoost | 20.10 | 95 | 13 | 54 | 1298 | 0.880 | 0.786–0.937 | 0.960 | 0.936–0.980 | 0.638 | 0.487–0.751 | 0.739 | 0.603–0.826 | 0.568 | 0.986 |
| LSTM | 19.80 | 85 | 23 | 43 | 1309 | 0.787 | 0.667–0.861 | 0.968 | 0.945–0.986 | 0.664 | 0.500–0.793 | 0.720 | 0.588–0.806 | 0.506 | 0.982 |

TP = true positive; FN = false negative; FP = false positive; TN = true negative; ROC = receiver operating characteristic; AUC = area under the receiver operating characteristic curve; CI = confidence interval. All models evaluated on 1,460 readings including 108 cold-stress readings at 6-hour lead time. Bootstrap CIs: 1,000 chronological resamples. <sup>a</sup> SHY = same-hour-yesterday naive baseline.

**Figure 7.**
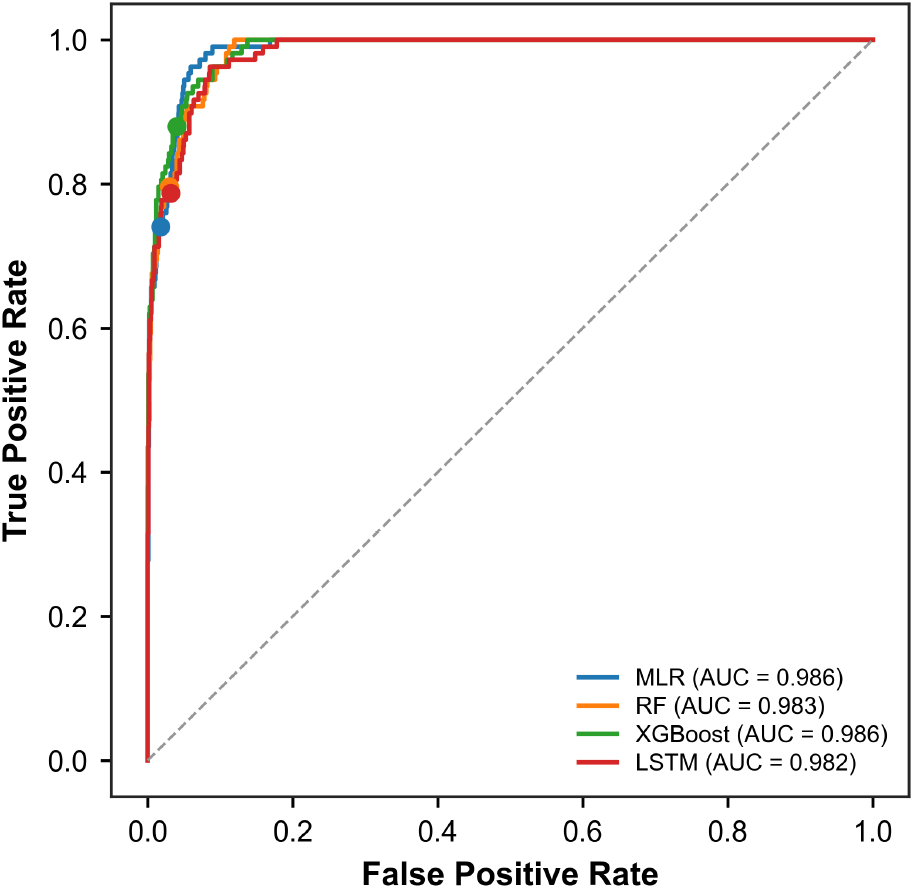
Receiver operating characteristic (ROC) curves for the four trained models evaluated on the independent 2025 test set for 6-hour-ahead cold-stress classification. Filled circles mark the validation-selected operating point for each model. Area under the curve (AUC) values are given in the legend.

**Figure 8.**
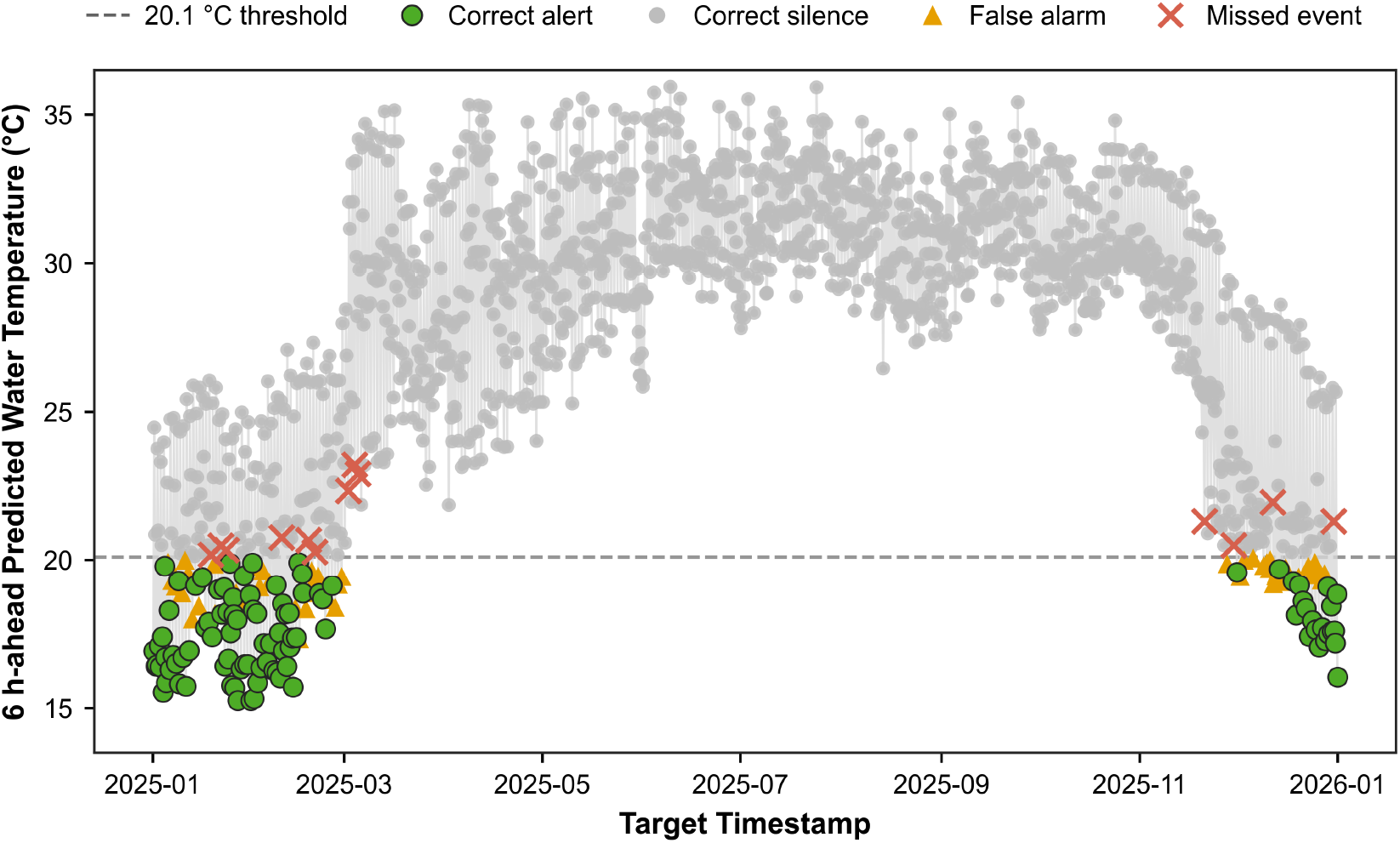
XGBoost 6-hour-ahead cold-stress warning output across the full independent 2025 test year. Observations are classified as correct alerts, false alarms, missed events, or correct silences. The dashed line marks the 20.1°C decision threshold.

Warning performance varied across daily target hours, as shown in Figure 9 for the 2024-25 winter season. The 06:00 target detected 73 of 74 cold-stress readings (98.6%) with 13 false alarms. The 00:00 target detected 45 of 49 cold-stress readings (91.8%) with 22 false alarms. The 18:00 target detected 8 of 14 cold-stress readings (57.1%) with 5 false alarms. The 12:00 target is excluded because no water temperature readings at that hour fell below 20°C in 2025.

**Figure 9.**
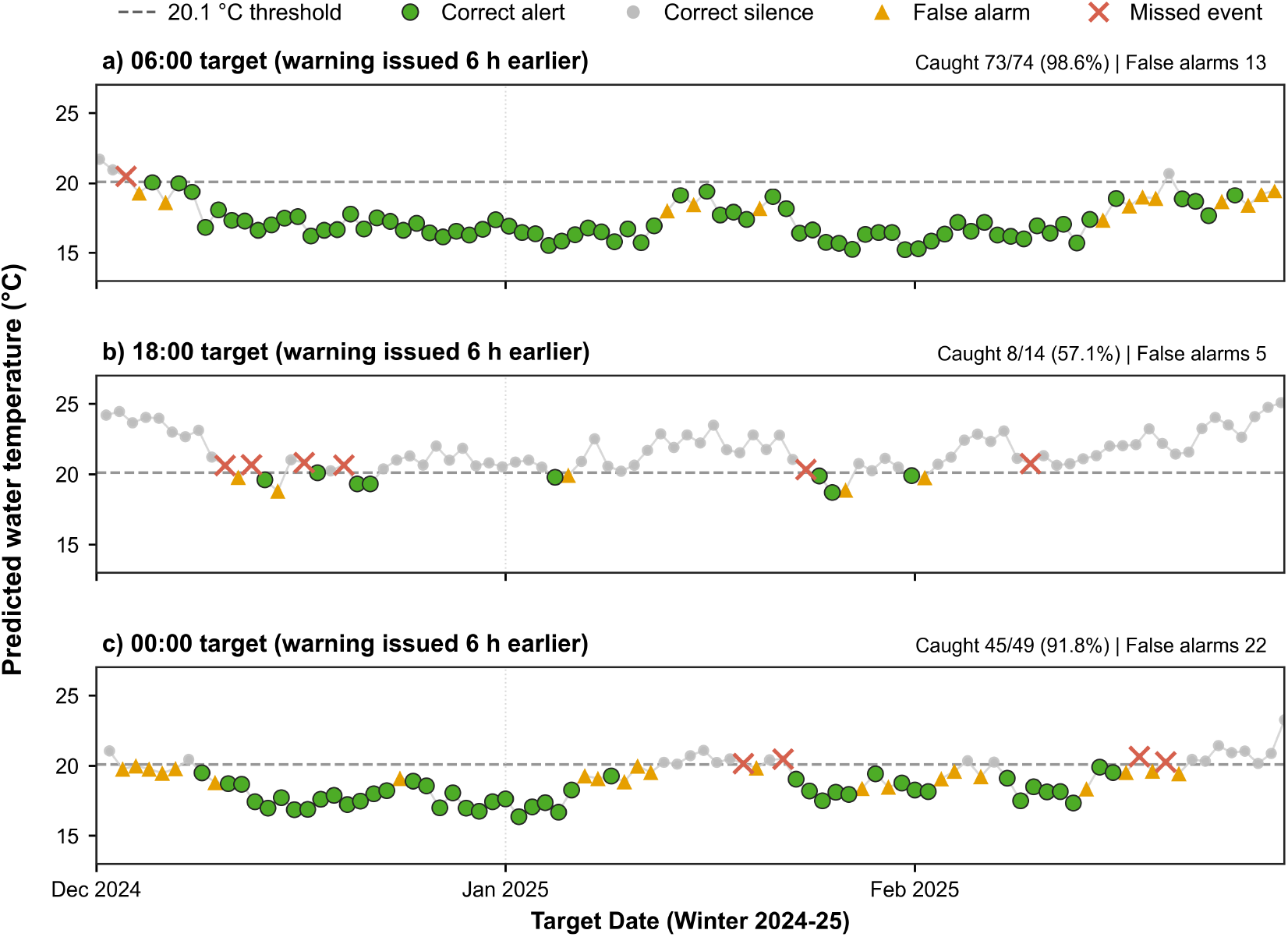
XGBoost 6-hour-ahead cold-stress warning output during the 2024–25 winter season at three daily target times: (a) 06:00, (b) 18:00, and (c) 00:00. Predicted water temperature is plotted against time with individual observations classified as correct alerts, false alarms, missed events, or correct silences; the dashed line marks the 20.1°C decision threshold.

Lead-time sensitivity analysis showed a non-monotonic trade-off for XGBoost (Figure 10; Table S4). At 6 h, sensitivity was 88.0%, precision was 63.8% and F1 was 73.9%. At 12 h, sensitivity declined to 76.9% while precision increased to 68.0% (F1 = 72.2%). At 18 h, the validation-optimized trigger increased to 21.1°C, sensitivity recovered to 86.1%, false alarms increased to 87 and precision declined to 51.7%. At 24 h, sensitivity was 80.6%, precision was 62.6% and F1 was 70.4%. The 6-h horizon produced the highest F1 across all tested lead times.

**Figure 10.**
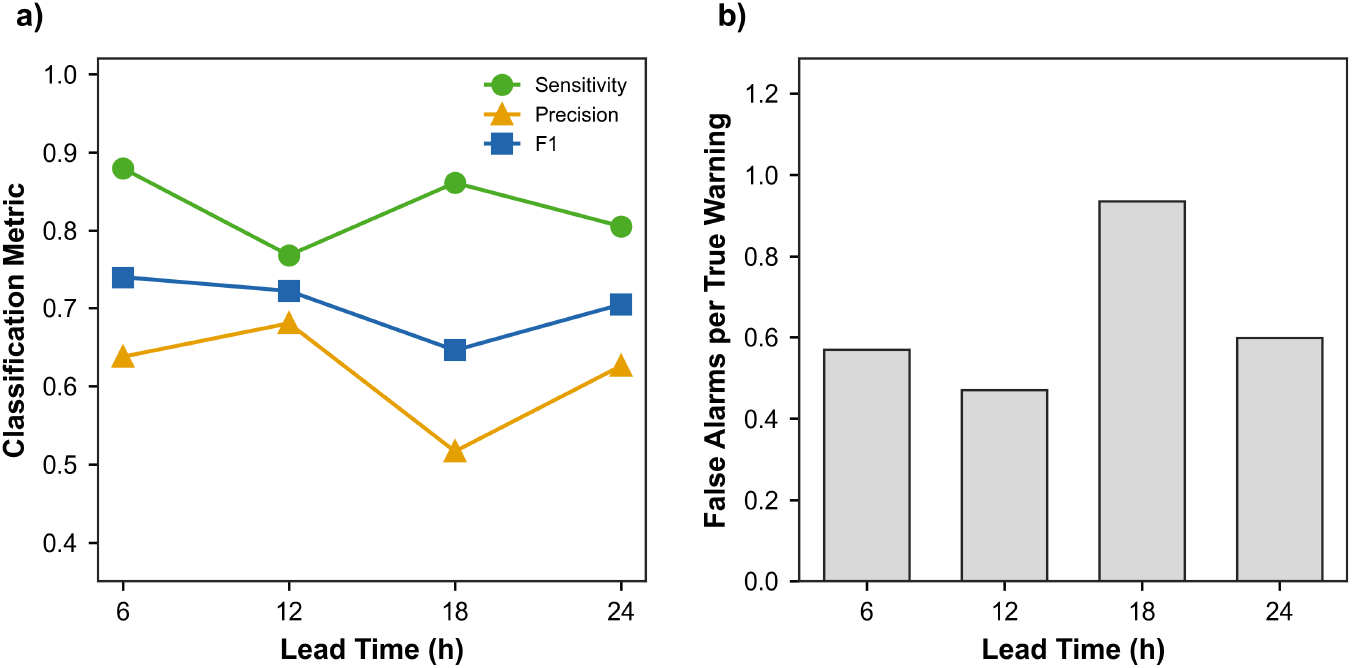
Cold-stress warning performance of XGBoost at forecast lead times of 6, 12, 18, and 24 hours. (a) Sensitivity, precision, and F1 score; (b) false alarms per true warning.

## Discussion

This study identifies winter cold stress as the primary thermal risk in the monitored nursery pond. As a warm-water species, Nile tilapia experiences reduced feeding, growth, and immunocompetence below approximately 20°C (Charo-Karisa et al., 2005; Nobrega et al., 2020). For nursery operations in particular, where small fish have limited energy reserves and high surface-area-to-volume ratios, sublethal cold exposure represents a more persistent management concern than episodic heat (Nobrega et al., 2020). The record contained 133 cold-stress days against only 4 heat-stress days (Tables 1 and 2; Figure 1), shifting management emphasis from annual temperature prediction to winter risk detection. For comparable low-resource hatchery and nursery settings, this finding reinforces the need for species-specific and context-specific climate advisories rather than generic weather warnings (Hossain et al., 2021; Montes et al., 2022).

The difference between 2024 and 2025 illustrates why multi-year records are needed for stable pond-level decision support (Table 2; Figure 3). The highest cold-stress burden occurred in 2024, whereas the independent test year, 2025, was milder. This contrast is methodologically useful: the model was trained on a period containing stronger cold exposure and tested on a year with fewer cold events. The dataset characterizes one pond across four winters; it is not a regional climate trend estimate.

The air-water relationship was strong over the full annual range but weaker in winter and within the 18-22°C boundary zone (Figure 4; Table S3). The weakening is not a physical break in pond thermodynamics at 20°C; it reflects the decision-zone limitation of air temperature as a proxy for pond-water temperature. The boundary-zone correlation drop is largely a statistical artefact of restricted-range correlation (Bland and Altman, 2011): when variance is truncated, the Pearson coefficient compresses even when the underlying slope remains stable. The boundary-zone slope of 0.775, close to the full-record slope of 0.922, confirms that the air-water thermodynamic link persists, but predictive uncertainty increases precisely where cold-stress classification is most sensitive. Seasonal diurnal profiles showed the expected thermal inertia signature: pond water was warmer than the air around dawn and cooler near midday in winter (Figure 2), a lag that a single same-time air reading cannot fully resolve. Similar lags have been documented in shallow aquatic systems (Caissie, 2006; Mohseni and Stefan, 1999) and represent a fundamental constraint on air-temperature-driven warning frameworks.

Within the boundary zone, solar radiation was lower on cold-stress days than on non-cold days (Cohen’s d = -0.39; Figure 5), consistent with Siddique et al. (2024), who identified solar intensity as an important factor for pond water temperature in a comparable Bangladesh tilapia pond culture system. Adding solar radiation to air temperature increased classification AUC only marginally, from 0.782 to 0.789, consistent with the full-record OLS result, where solar radiation was not significant once air temperature was accounted for. This also fits evidence that radiative forcing in shallow aquaculture ponds is secondary to convective and sensible heat exchange at sub-daily timescales (Caissie, 2006; Resnick et al., 2024). The NASA POWER data provide gridded regional estimates rather than site-specific pond measurements, and the limited AUC gain may partly reflect spatial averaging. In practice, the framework does not require solar radiation inputs, which is advantageous for settings where only a thermometer is available.

In 6-h regression, the same-hour-yesterday baseline (MAE = 1.117°C) nearly matched XGBoost (MAE = 1.116°C), confirming that shallow-pond thermal memory at the same clock hour is the dominant short-lead signal (Table 3). This is consistent with Zambrano et al. (2021), who showed that random forests can accurately forecast pond water-quality variables from as few as two manual measurements per day. The value of trained models is not that they outperform this naive signal on aggregate accuracy, but that they can be tuned into biologically relevant decision thresholds. RF achieved the smallest cold-zone bias (+0.30°C), but XGBoost combined a comparably low bias (+0.36°C) with the smallest train-test MAE gap and the highest warning sensitivity (Table 4), making it the model carried forward for the warning analysis. We had initially expected LSTM to perform better than tree-based models given its sequence-awareness, but the compact 6-hour lag structure may not provide enough temporal depth for recurrent architectures to demonstrate their advantage. This is consistent with findings from short-horizon environmental temperature forecasting where gradient-boosted trees outperform recurrent networks when the lag structure is compact (Feigl et al., 2021; Huan et al., 2020; Ren et al., 2020). The larger generalization gap for RF relative to XGBoost reflects the different regularization properties of bagging and boosting ensembles (Breiman, 2001; Chen and Guestrin, 2016). The milder 2025 cold season reduced aggregate test difficulty for MLR and LSTM relative to the fitting period (Table 2), explaining their negative train-test gaps.

The warning results are best understood as a threshold decision problem rather than a single regression ranking (Table 4). MLR provided the most balanced classifier by F1 and precision, whereas XGBoost achieved higher sensitivity (0.880) at a higher false-alarm cost, representing the more protective operating point. Because cold-stress readings were the minority class, ROC AUC alone is insufficient for model selection; precision, F1, and the false-alarm burden must be evaluated jointly (Saito and Rehmsmeier, 2015). The bootstrap intervals in Table 4 overlap substantially across the top models and the same-hour-yesterday baseline, so these models are not statistically distinguishable at this sample size. The choice between them should therefore be guided by farm-specific tolerance for missed events versus unnecessary alerts rather than by a metric difference the data cannot resolve. Alerts that trigger low-cost responses, such as closer observation, delayed feeding or postponed transfer, carry a different risk profile from those that trigger expensive or irreversible intervention.

Lead-time analysis showed that longer XGBoost horizons did not improve on the 6-h F1 (Figure 10; Table S4). The 12-h horizon traded sensitivity for precision; the 18-h horizon recovered sensitivity at the cost of substantially more false alarms; and the 24-h horizon retained useful discrimination but did not exceed the 6-h operating point. These results support a tiered framework in which 6-h alerts serve as the primary operational warning and 12-24 h outputs are treated as planning-level advisories.

The proposed framework addresses a distinct operating case from sensor-heavy smart aquaculture systems. Reviews of aquaculture monitoring describe the value of IoT networks, biosensors, and cloud dashboards for continuous data streams (Su et al., 2020; Zhao et al., 2021), and recent fishpond cold-snap systems combine numerical weather models with in-pond sensors for real-time warnings (Li et al., 2024). These systems are effective where infrastructure exists but do not match all farm contexts. The present study demonstrates that a warning system built from routine manual thermometer readings can achieve meaningful cold-stress detection at 6-h lead time, converting a practice that farmers already perform into a decision-support tool without requiring capital investment in automated hardware. This positioning is relevant for small-holder nursery operations where sensor infrastructure remains limited (Hossain et al., 2021) and aligns with broader efforts to make ML-based aquaculture tools accessible in data-scarce settings (Zambrano et al., 2021).

Our results support all four hypotheses. Cold stress was the dominant thermal hazard and varied substantially across years, with the heaviest burden in 2024 (H1). Solar radiation was associated with cold-stress status within the boundary zone but added little classification value beyond air temperature (H2). Nonlinear models reduced cold-zone bias relative to MLR, most clearly for XGBoost and RF, though the full-test MAE differences were small (H3). At 6-h lead time, the thermometer-based warning framework delivered actionable skill in the independent 2025 test year (H4). These findings have three practical implications for aquaculture management. First, the warning framework does not require solar radiation data. Second, nonlinear modelling offers modest but real benefits for cold-zone prediction. Third, thermometer-based warnings can provide operational value where IoT infrastructure is unavailable. The study was conducted on a single nursery pond over four winters; multi-pond validation across contrasting pond types and cold-season intensities is the necessary next step.

## Conclusions

This study developed and tested a cold-stress early warning framework for aquaculture nursery ponds using manual thermometer readings collected at 6-hour intervals. The framework was evaluated on a Nile tilapia nursery pond in Bangladesh using four years of field data, with the final year held out as an independent test. Same-target-hour thermal memory was a strong predictor, but trained models achieved better precision-sensitivity tradeoffs at the warning stage, reducing false alarms while maintaining detection rates. Among trained models, XGBoost provided the more protective alert and MLR the most balanced classification. The 6-hour lead time produced the best warning performance, supporting a tiered approach where 6-hour alerts serve as operational warnings and longer horizons serve as planning advisories. Future work should validate the framework across ponds of varying size, depth, and management practice, across other cold-sensitive species, and in comparable settings where automated monitoring remains impractical.

## DATA AVAILABILITY

Field temperature data supporting the findings of this study are available from the corresponding author upon reasonable request. NASA POWER data are publicly available at https://power.larc.nasa.gov/.

## CODE AVAILABILITY

The analysis code and reproducible workflow are available in a public GitHub repository (https://github.com/imranbinyounos/cold-stress-early-warning-aquaculture).

## AUTHOR CONTRIBUTIONS

Imran Bin Younos: Conceptualization, Data curation, Formal analysis, Methodology, Software, Visualization, Writing – Original Draft. Nushrat Jahan: Methodology, Validation, Writing – Review & Editing.

## ACKNOWLEDGEMENTS

The authors gratefully acknowledge the hatchery personnel for their assistance with routine pond management, manual temperature monitoring, and maintenance of field records during the study period.

