## Supplementary Information for "A Low-Resource Machine-Learning Framework for Cold-Stress Early Warning in Aquaculture Nursery Ponds Using Manual Temperature Readings"

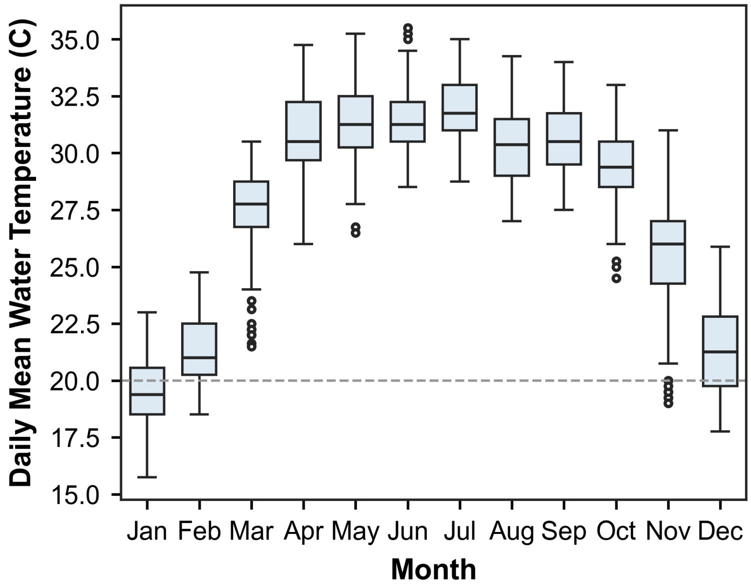


**Figure S1.** Monthly distributions of daily mean pond water temperature pooled across all four study years (2022–2025). The dashed line at 20 °C indicates the cold-stress threshold.

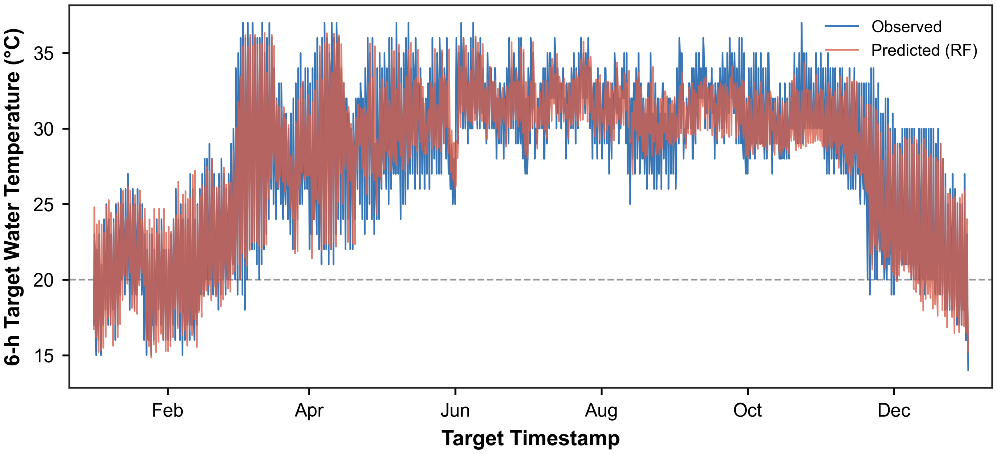


**Figure S2.** Observed and random forest-predicted six-hour-ahead pond water temperature across the 2025 independent test year. The dashed line at 20°C indicates the cold-stress threshold.

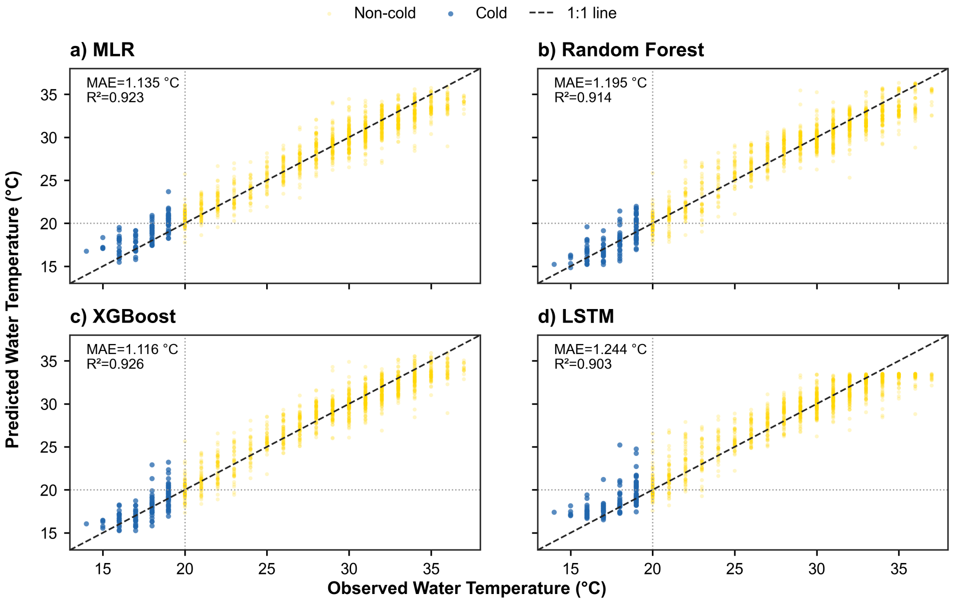


**Figure S3.** Observed versus predicted six-hour-ahead pond water temperature on the 2025 test set for (a) MLR, (b) random forest, (c) XGBoost, and (d) LSTM. Cold observations (below 20°C) are shown in blue; the dashed diagonal represents perfect agreement. MAE and $R^2$ are annotated within each panel.

**Table S1.** Seasonal six-hour-ahead regression error metrics for each trained model on the 2025 test year.

| Model | Season | n | MAE (°C) | RMSE (°C) | $R^2$ | Bias (°C) |
|---|---|---|---|---|---|---|
| MLR | Winter | 360 | 1.047 | 1.305 | 0.866 | 0.226 |
| | Pre-monsoon | 368 | 1.273 | 1.612 | 0.856 | -0.310 |
| | Monsoon | 488 | 1.169 | 1.485 | 0.584 | -0.184 |
| | Post-monsoon | 244 | 0.987 | 1.337 | 0.866 | -0.031 |
| RF | Winter | 360 | 1.016 | 1.272 | 0.873 | -0.211 |
| | Pre-monsoon | 368 | 1.356 | 1.768 | 0.827 | 0.108 |
| | Monsoon | 488 | 1.176 | 1.508 | 0.572 | -0.166 |
| | Post-monsoon | 244 | 1.252 | 1.570 | 0.815 | -0.331 |
| XGBoost | Winter | 360 | 1.020 | 1.256 | 0.876 | -0.221 |
| | Pre-monsoon | 368 | 1.294 | 1.669 | 0.846 | 0.064 |
| | Monsoon | 488 | 1.094 | 1.384 | 0.639 | -0.175 |
| | Post-monsoon | 244 | 1.034 | 1.349 | 0.863 | -0.142 |
| LSTM | Winter | 360 | 1.014 | 1.289 | 0.869 | 0.179 |
| | Pre-monsoon | 368 | 1.698 | 2.186 | 0.735 | 0.138 |
| | Monsoon | 488 | 1.148 | 1.462 | 0.597 | -0.079 |
| | Post-monsoon | 244 | 1.092 | 1.413 | 0.850 | 0.005 |

MAE = mean absolute error; RMSE = root mean square error. Evaluated on the 2025 independent test year.

**Table S2.** Cold-stress warning performance by time of day on the independent 2025 test year.

| Model | Decision hour | Target hour | Lead (h) | Trigger (°C) | TP | FN | FP | TN | Sensitivity | Specificity | Precision | F1 | False alarms per true warning | ROC AUC | Observed cold readings |
|---|---|---|---|---|---|---|---|---|---|---|---|---|---|---|---|
| Persistence | 0 | 6 | 6 | 21.10 | 58 | 6 | 29 | 272 | 0.906 | 0.904 | 0.667 | 0.768 | 0.500 | 0.948 | 64 |
| | 6 | 12 | 6 | 21.10 | 0 | 0 | 100 | 265 | 0.000 | 0.726 | – | – | – | – | 0 |
| | 12 | 18 | 6 | 21.10 | 1 | 7 | 0 | 357 | 0.125 | 1.000 | 1.000 | 0.222 | 0.000 | 0.969 | 8 |
| | 18 | 0 | 6 | 21.10 | 28 | 8 | 22 | 307 | 0.778 | 0.933 | 0.560 | 0.651 | 0.786 | 0.955 | 36 |
| SHY[a] | 0 | 6 | 6 | 20.10 | 64 | 0 | 24 | 277 | 1.000 | 0.920 | 0.727 | 0.842 | 0.375 | 0.989 | 64 |
| | 6 | 12 | 6 | 20.10 | 0 | 0 | 0 | 365 | 0.000 | 1.000 | – | – | – | – | 0 |
| | 12 | 18 | 6 | 20.10 | 8 | 0 | 16 | 341 | 1.000 | 0.955 | 0.333 | 0.500 | 2.000 | 0.981 | 8 |
| | 18 | 0 | 6 | 20.10 | 31 | 5 | 34 | 295 | 0.861 | 0.897 | 0.477 | 0.614 | 1.097 | 0.958 | 36 |
| MLR | 0 | 6 | 6 | 20.10 | 52 | 12 | 7 | 294 | 0.812 | 0.977 | 0.881 | 0.846 | 0.135 | 0.989 | 64 |
| | 6 | 12 | 6 | 20.10 | 0 | 0 | 0 | 365 | 0.000 | 1.000 | – | – | – | – | 0 |
| | 12 | 18 | 6 | 20.10 | 4 | 4 | 2 | 355 | 0.500 | 0.994 | 0.667 | 0.571 | 0.500 | 0.986 | 8 |
| | 18 | 0 | 6 | 20.10 | 24 | 12 | 15 | 314 | 0.667 | 0.954 | 0.615 | 0.640 | 0.625 | 0.962 | 36 |
| RF | 0 | 6 | 6 | 19.60 | 57 | 7 | 16 | 285 | 0.891 | 0.947 | 0.781 | 0.832 | 0.281 | 0.985 | 64 |
| | 6 | 12 | 6 | 19.60 | 0 | 0 | 0 | 365 | 0.000 | 1.000 | – | – | – | – | 0 |
| | 12 | 18 | 6 | 19.60 | 3 | 5 | 3 | 354 | 0.375 | 0.992 | 0.500 | 0.429 | 1.000 | 0.977 | 8 |
| | 18 | 0 | 6 | 19.60 | 26 | 10 | 21 | 308 | 0.722 | 0.936 | 0.553 | 0.627 | 0.808 | 0.957 | 36 |
| XGBoost | 0 | 6 | 6 | 20.10 | 61 | 3 | 20 | 281 | 0.953 | 0.934 | 0.753 | 0.841 | 0.328 | 0.989 | 64 |
| | 6 | 12 | 6 | 20.10 | 0 | 0 | 0 | 365 | 0.000 | 1.000 | – | – | – | – | 0 |
| | 12 | 18 | 6 | 20.10 | 5 | 3 | 3 | 354 | 0.625 | 0.992 | 0.625 | 0.625 | 0.600 | 0.978 | 8 |
| | 18 | 0 | 6 | 20.10 | 29 | 7 | 31 | 298 | 0.806 | 0.906 | 0.483 | 0.604 | 1.069 | 0.962 | 36 |
| LSTM | 0 | 6 | 6 | 19.80 | 57 | 7 | 16 | 285 | 0.891 | 0.947 | 0.781 | 0.832 | 0.281 | 0.985 | 64 |
| | 6 | 12 | 6 | 19.80 | 0 | 0 | 0 | 365 | 0.000 | 1.000 | – | – | – | – | 0 |
| | 12 | 18 | 6 | 19.80 | 1 | 7 | 1 | 356 | 0.125 | 0.997 | 0.500 | 0.200 | 1.000 | 0.955 | 8 |
| | 18 | 0 | 6 | 19.80 | 27 | 9 | 26 | 303 | 0.750 | 0.921 | 0.509 | 0.607 | 0.963 | 0.960 | 36 |

TP = true positive; FN = false negative; FP = false positive; TN = true negative; ROC = receiver operating characteristic; AUC = area under the receiver operating characteristic curve; [a] SHY = same-hour-yesterday naive baseline.

**Table S3.** Air-water temperature coupling statistics for the full four-year record, by season, and within the 18–22°C air-temperature boundary zone.

| Group | n | Pearson r | Pearson p | Slope | Intercept | $R^2$ |
|---|---|---|---|---|---|---|
| Full record | 1461 | 0.976 | <0.001 | 0.922 | 2.318 | 0.952 |
| Monsoon | 488 | 0.860 | <0.001 | 0.782 | 6.895 | 0.740 |
| Post-monsoon | 244 | 0.951 | <0.001 | 0.888 | 3.224 | 0.904 |
| Pre-monsoon | 368 | 0.953 | <0.001 | 0.729 | 8.093 | 0.909 |
| Winter | 361 | 0.776 | <0.001 | 0.751 | 5.455 | 0.603 |
| 18–22°C boundary zone | 295 | 0.541 | <0.001 | 0.775 | 5.015 | 0.292 |

OLS = ordinary least squares regression (daily mean air temperature as predictor; pond water temperature as response). All $p < 0.001$.

**Table S4.** XGBoost regression accuracy and cold-stress warning performance across forecast lead times of 6 to 24 hours on the independent 2025 test set.

| Lead time (h) | Trigger (°C) | Test MAE (°C) | Test RMSE (°C) | Test $R^2$ | TP | FN | FP | TN | Sensitivity | Specificity | Precision | F1 | ROC AUC | False alarms per true warning |
|---|---|---|---|---|---|---|---|---|---|---|---|---|---|---|
| 6 | 20.10 | 1.116 | 1.427 | 0.926 | 95 | 13 | 54 | 1298 | 0.880 | 0.960 | 0.638 | 0.739 | 0.986 | 0.568 |
| 12 | 20.20 | 1.194 | 1.530 | 0.915 | 83 | 25 | 39 | 1313 | 0.769 | 0.971 | 0.680 | 0.722 | 0.982 | 0.470 |
| 18 | 21.10 | 1.190 | 1.528 | 0.915 | 93 | 15 | 87 | 1265 | 0.861 | 0.936 | 0.517 | 0.646 | 0.980 | 0.935 |
| 24 | 20.20 | 1.188 | 1.521 | 0.916 | 87 | 21 | 52 | 1300 | 0.806 | 0.962 | 0.626 | 0.704 | 0.981 | 0.598 |

MAE = mean absolute error; RMSE = root mean square error; AUC = area under the receiver operating characteristic curve; TP = true positive; FN = false negative; FP = false positive; TN = true negative. Separate XGBoost models fitted per horizon. Evaluated on 1,460 readings (108 cold-stress).
